# The Intrinsic Manifold of Spontaneous Activity Constrains Cortical Responses to Naturalistic Stimuli

**DOI:** 10.64898/2026.02.21.707183

**Authors:** Jung-Hoon Kim, Yizhen Zhang, Owen MacKenzie, Xiaokai Wang, David Brang, Scott J. Peltier, Zhongming Liu

## Abstract

The cerebral cortex constrains its spontaneous activity to a low-dimensional manifold learnable from resting state functional magnetic resonance imaging (fMRI) data. However, it remains unclear whether this intrinsic manifold also captures cortical responses to complex, naturalistic stimuli. To test this, we pretrained a deep variational autoencoder model on 3-Tesla resting-state fMRI data to learn the latent structure of spontaneous activity, and then applied this model without finetuning to 7-Tesla fMRI data acquired during movie-watching. Despite the different field strengths, the model generalized robustly from resting to movie-watching states. The latent representation of stimulus-evoked responses was confined to a subspace that occupied about 13% of the latent space spanned by spontaneous activity, demonstrating that task-related neural responses do not require a distinct representational space. By representing cortical dynamics as an evolving latent trajectory, we found striking differences across individuals or between brain states. During movie watching, the velocity of the latent trajectory provided a reliable marker of cortical engagement, and its temporal structure was highly reliable and sensitive to naturalistic events. These findings suggest that the intrinsic manifold of spontaneous activity forms a full reservoir of cortical states that the brain can differentially engage when interacting with the external environment.

## Introduction

The cerebral cortex is a dynamical system, composed of billions of neurons, thousands of voxels, and hundreds of regions. Despite this vast dimensionality, cortical activity does not wander randomly through all possible configurations. Instead, it is intrinsically constrained by structural connectivity and cortical folding patterns (Honey et al., 2009; Pang et al., 2023), and driven by a variety of physiological and neural sources (Lu et al., 2019). These constraints shape neural information flow across circuits and networks, induce complex and nonlinear interactions, and confine brain states to a functional reservoir that is rich yet bounded (Deco et al., 2013; Breakspear, 2017).

This bounded reservoir constitutes a “manifold” (Gallego et al., 2017). Formally, this manifold is a low-dimensional, nonlinear, and closed surface embedded within the high-dimensional space spanned by *N* voxels. Dynamic cortical activity resides on and evolves along this manifold. Its trajectory is constrained by a limited set of local directions (or “intrinsic dimensions”) which define the valid paths of movement everywhere on the manifold. By manifold learning, we can explicitly model this manifold and define a low-dimensional “latent space” (Pandarinath et al., 2018; Schneider et al., 2023), where each latent variable captures a unique intrinsic dimension (Figure 1).

**Figure 1:**
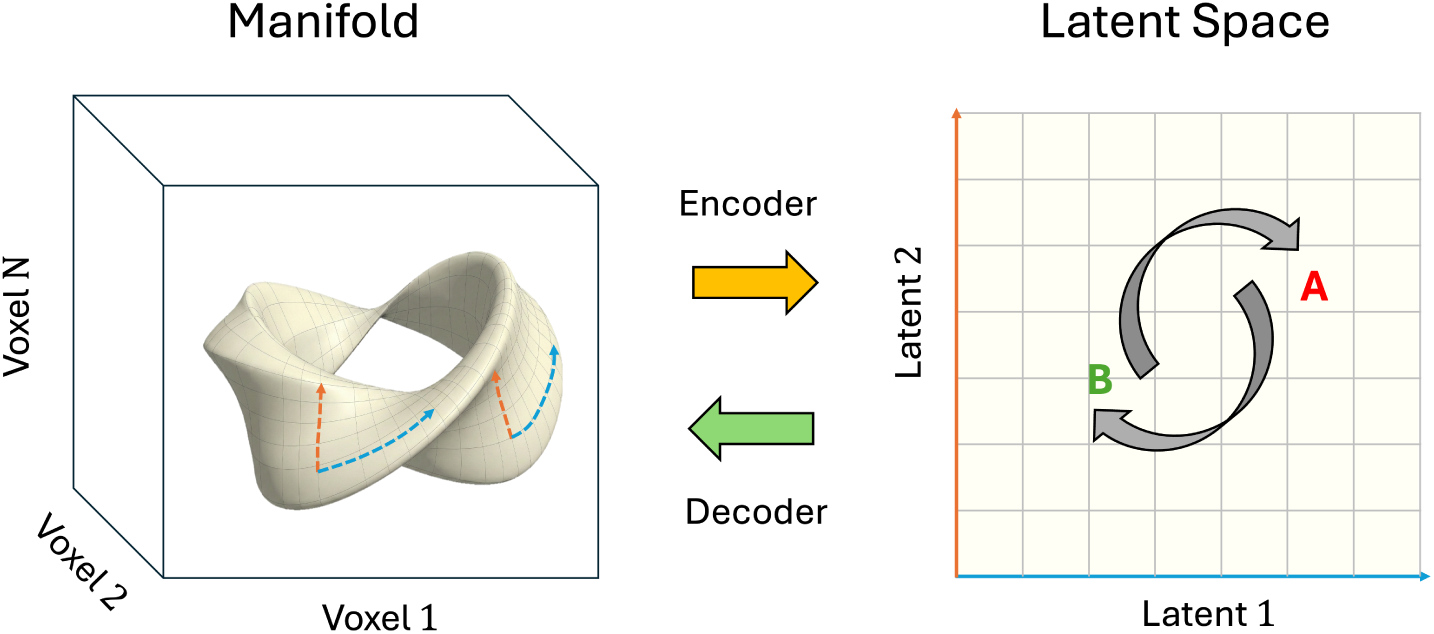
Manifold embedding of cortical dynamics. (Left) Cortical activity is represented in a high-dimensional space spanned by *N* voxels. The signal dynamics are constrained to a nonlinear manifold (surface) embedded within the voxel space. The dashed arrows (blue and orange) illustrate the manifold’s intrinsic dimensions. (Right) An encoder maps the nonlinear manifold onto a low-dimensional latent space, while the decoder performs the reverse mapping. The axes Latent 1 (blue) and Latent 2 (orange) correspond to the intrinsic dimensions of the manifold. The curved arrows illustrate latent trajectories that lead the transition from one state (denoted as **A**) to another (denoted as **B**).

To learn the manifold of cortical activity, it is desirable to use training data that sufficiently samples the underlying distribution. Spontaneous activity is well suited for this purpose, as the cortex exhibits rich, diverse, and organized activity patterns in the absence of any overt tasks (Biswal et al., 1995; Fox & Raichle, 2007; Greicius et al., 2003). In a mesoscopic scale, spontaneous activity in the visual cortex cycles through spatial maps that precisely match those evoked by visual stimulation (Kenet et al., 2003). In the whole-brain scale, resting state functional magnetic resonance imaging (rs-fMRI) studies have established the correspondence between spontaneous and task-evoked co-activation patterns (Smith et al., 2009; Cole et al., 2014; Tavor et al., 2016). In addition, rs-fMRI data are abundant and publicly available, e.g., the Human Connectome Project (HCP) (Van Essen et al., 2013), making it feasible to train deep models.

Building upon this conceptual framework, our prior work demonstrated that a deep variational autoencoder (VAE) (Kingma & Welling, 2013; Higgins et al., 2017) trained solely on rs-fMRI data can model the intrinsic manifold of spontaneous cortical activity and learn the associated latent space (Kim et al., 2021). However, it remains unclear whether this intrinsic manifold of task-free activity generalizes to complex, dynamic cortical activity during tasks. To test this, we applied our pretrained VAE model to fMRI data acquired during movie watching. We sought to determine whether the rich cortical responses to naturalistic stimuli are confined to the intrinsic manifold defined by spontaneous activity, and whether the latent trajectories can effectively capture the attributes and structure of naturalistic experiences.

## Methods and Materials

### VAE pretrained with 3-T rs-fMRI data

Our VAE model was pretrained (Kim et al., 2021) based on 3-T rs-fMRI data released from HCP (Van Essen et al., 2013). Briefly, the model employs an encoder-decoder architecture (Figure 2a). The encoder consists of five convolutional layers and one fully connected layer, transforming an instantaneous pattern of cortical activity into 256 latent variables, each modeled as a Gaussian distribution. The decoder consists of one fully connected layer and five transposed convolutional layers, reconstructing or generating the input from samples of the latent distribution. The learning objective during pretraining is to reconstruct the input while enforcing the latent variables to be close to independent standard normal distributions. During training or testing, the model’s input and output consist of two pixel arrays, representing the left and right cortical hemispheres, inflated to spheres and evenly resampled to 192 *×* 192 grids defined on the basis of azimuth and sin(elevation). See details in Kim et al. (2021).

**Figure 2:**
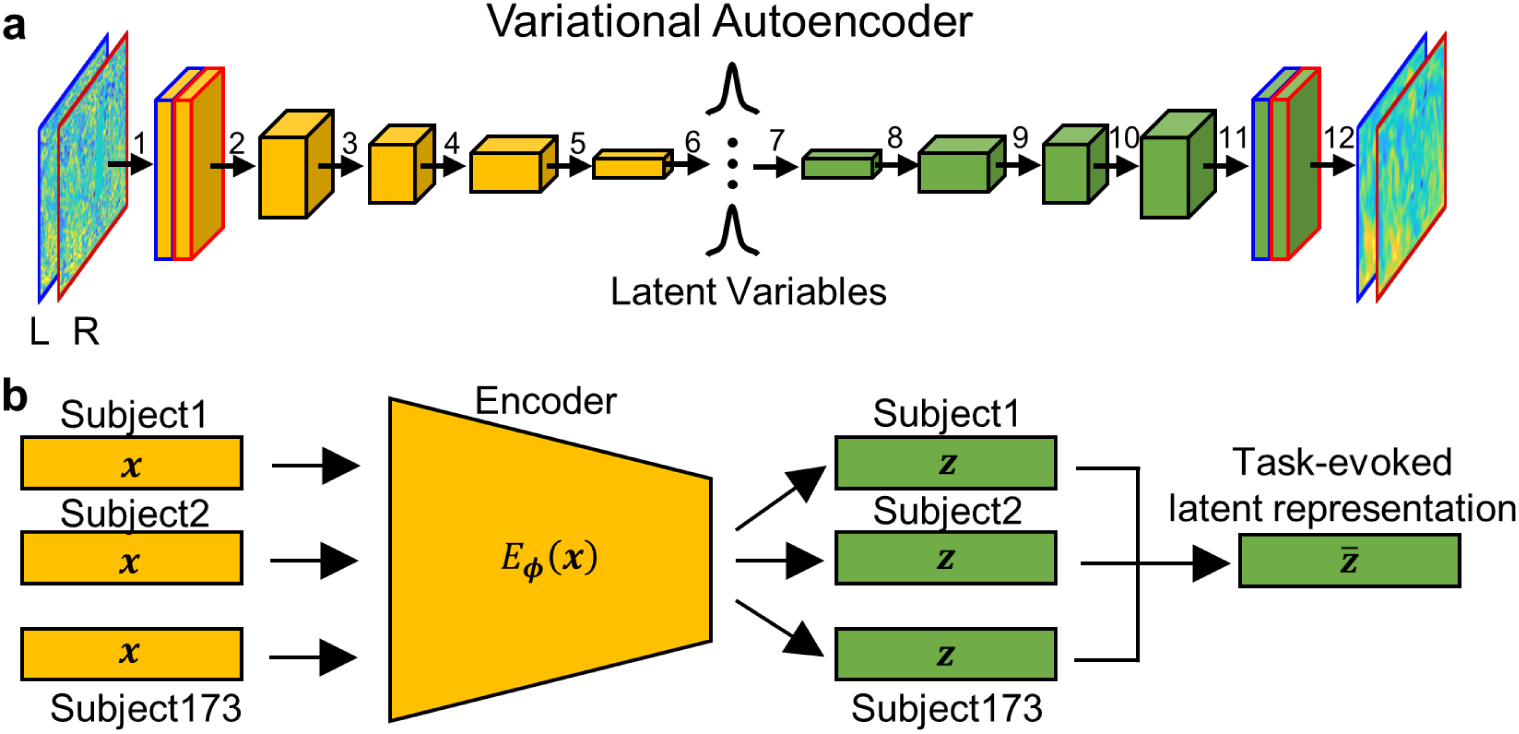
VAE architecture and inference of task fMRI. **a**. The VAE architecture consists of the encoder (yellow) and the decoder (green). Numbered arrows indicate specific layer operations: 1: Convolution (kernel size=8, stride=2, padding=3) with ReLU activation; 2–5: Convolution (kernel size=4, stride=2, padding=1) with ReLU activation; 6: Fully connected layer with reparameterization; 7: Fully connected layer with ReLU activation; 8–11: Transposed convolution (kernel size=4, stride=2, padding=1) with ReLU activation; 12: Transposed convolution (kernel size=8, stride=2, padding=3). Blue and red frames represent the input/output images for the left and right hemispheres, respectively. **b**. Extraction of task-evoked latent representations. The pretrained encoder (*E_ϕ_*) maps individual fMRI data (*x*) to subject-specific latent representations (*z*). The group-level task-evoked representation (*z̄*) is computed by averaging these latent vectors across all 173 subjects to isolate stimulus-driven activity.

### 7-T rs-fMRI and movie-fMRI data

We used the pretrained VAE model to analyze resting-state and task-based fMRI data from 173 young adults from the HCP 7-T data release (Van Essen et al., 2013; Uğurbil et al., 2013). Each subject underwent multiple fMRI sessions across different days using resting-state or movie-watching paradigms. Each resting-state session included 900 time points (15 minutes). Movie-watching sessions were at least 15 minutes each with slight variations due to differing movie durations: sessions 1 to 4 had 921, 918, 915, and 901 time points, respectively.

As detailed in Uğurbil et al. (2013), data were acquired using a gradient-echo echo-planar imaging (EPI) sequence with the following parameters: repetition time (TR) = 1000 ms, echo time = 22.2 ms, flip angle = 45*^◦^*, field of view = 208 *×* 208 mm, matrix size = 130 *×* 130, voxel size = 1.6 mm isotropic, 85 slices, multiband factor = 5, in-plane acceleration = 2, partial Fourier sampling = 7*/*8, echo spacing = 0.64 ms, and bandwidth = 1924 Hz/Px. Data were preprocessed using the minimal preprocessing pipeline (Glasser et al., 2013), ICA-FIX (Griffanti et al., 2014; Salimi-Khorshidi et al., 2014), temporal detrending (by removing third-order polynomials), bandpass filtering (0.01 *−* 0.1 Hz), and normalization (to zero mean and unit variance) for each voxel.

All subjects watched the same videos across four sessions. Sessions 1 and 3 included short video clips available under Creative Commons licenses on Vimeo, while sessions 2 and 4 included segments from Hollywood movies. Details of the video clips are provided in Supplementary Table 1. At the end of each session, an 83-second video clip was presented for test-retest across sessions. Each clip began and ended with a 20-second rest period displaying “REST” on a black screen. See Appendix Table S1 for the information about the video fMRI data used in this study.

### Generalizability to 7-T and Task-Evoked Data

To assess its generalizability, we applied the VAE model (pretrained on 3-T rs-fMRI data) directly to the 7-T rs-fMRI and movie-fMRI datasets without fine-tuning. The model encoded the input 7-T cortical patterns into latent vectors and reconstructed the inputs from these latent representations. We evaluated the reconstruction accuracy by computing the Pearson correlation between the reconstructed and the input patterns after spatial smoothing by a 2-D Gaussian kernel with a full-width-half-maximum of 6 mm.

We compared the VAE’s performance with ICA implemented in FSL (Smith et al., 2004). We applied ICA to the same 3-T rs-fMRI training data to obtain 256 independent components, providing a linear latent space with the same dimension as the latent space obtained with VAE. These components were then used to represent and reconstruct the 7-T testing data, similar to the VAE approach. As a null baseline, we generated cortical patterns from randomly sampled latent variables (drawn from a standard Gaussian distribution) and assessed the reconstruction accuracy between these randomly generated patterns and the input patterns.

### Latent Space Visualization of State Transitions

We examined how transitions between resting and movie-watching states manifested in the latent space. The fMRI data from all subjects were processed through the pretrained VAE to obtain latent representations. We used t-distributed Stochastic Neighbor Embedding (t-SNE) (van der Maaten & Hinton, 2008) to visualize the latent representations from all subjects in a 2-D space. For demonstration, we focused on a 30-second period around the transition from rest to watching a specific video clip (Northwest). Specifically, the first 15 seconds (labeled as 0 to 15 seconds) were immediately prior to the movie onset, and the second 15 seconds (labeled as 15 to 30 seconds) were immediately after the movie onset. For visual inspection, the latent representations were color-coded either by time to highlight brain state transitions or by subject to highlight individual variations.

### Geometry and Dimensionality of the Task Evoked Subspace

To characterize task-evoked latent representations, we processed the movie-fMRI data from all subjects through the pretrained VAE and averaged the latent representations across subjects, isolating task-related activity (Figure 2b). Based on the task-evoked latent representations, we computed the covariance matrix, where each of-diagonal element measured the covariance between a pair of latent variables and each diagonal element measured the variance of a single latent variable. Principal component analysis (PCA) was applied to the covariance matrix to determine the number of principal components required to explain 90% of the variance. These components serve as orthogonal basis vectors, spanning a subspace with dimensionality defined by the number of principal components. We performed this analysis on latent representations from both the task-evoked and resting-state fMRI data for dimensionality comparison.

### Quantification of Latent Dynamics

We analyzed the latent trajectory during movie watching by computing the difference between consecutive latent representations, estimating the instantaneous velocity of the latent trajectory. The magnitude of this velocity quantified the rate of change at each time point, and its average over time within each movie quantified that movie’s overall level of cortical engagement.

We also examined the temporal characteristics of the latent velocity during inter-movie rest periods, each lasting 20 seconds. We extracted a 30-second time series of the latent velocity spanning 5 seconds before the onset and 5 seconds beyond the end of each 20-second inter-movie period. We averaged such time series across all subjects to obtain an aggregated latent response time-locked to the inter-movie period.

### Test-Retest Reliability and Inter-Subject Consistency

We evaluated the reliability of the latent trajectory across repeated presentations of the same movie (“Test-retest” movie). For each session, we computed the top 24 principal components (basis vectors) of the latent trajectory and projected the session-specific trajectory onto these components to obtain their corresponding dynamics. We assessed the similarity of these components and their dynamics across sessions using Pearson’s correlation coefficients.

We also evaluated the inter-subject consistency of the latent trajectory. First, we identified the shared dynamic patterns by performing PCA on the group-averaged latent trajectory, yielding a set of group-level principal components (basis vectors). To determine how well each individual expressed these shared patterns, we projected each subject’s latent trajectory onto these group-level principal components. We then calculated the correlation between each subject’s projected dynamics and the group-average dynamics. Finally, these correlation coefficients were Fisher z-transformed and averaged across all subjects to quantify the cross-subject consistency at the group level.

### Statistical Analysis

Statistical tests used here were implemented using MATLAB functions, including one-sample or paired-sample t-tests (ttest.m), two-sample t-test (ttest2.m), one-way analysis of variance (ANOVA) (anova1.m), and two-sample F-test (vartest2.m). Specifically, the statistical significance of any change in the latent velocity over different movie clips was evaluated using one-way ANOVA. Post hoc comparison between repetitions was conducted using the two-sample t-test. Multiple comparison correction was conducted by controlling the false-discovery rate (FDR) *q <* 0.05. For the velocity of the latent trajectory, the significance of response amplitudes (against 0) at each time point was evaluated using t-tests with FDR correction (*q <* 0.05).

## Results

We tested the generalizability of our VAE model trained on 3-T rs-fMRI data (Kim et al., 2021) to unseen, out-of-distribution fMRI data collected at different field strengths and under different brain states (i.e., watching movies). Using the HCP 7-T fMRI datasets (Van Essen et al., 2013; Uğurbil et al., 2013), we investigated the distribution and trajectory of latent representations of cortical activity during rest and movie watching and characterized the key distinctions in their engagement of cortical networks and dynamics. In addition, we investigated various factors that shaped the latent trajectory during movie watching, including audiovisual events, individual consistency and variation, and test-retest reliability.

### Generalizability from 3-T to 7-T and from Rest to Tasks

We tested our model’s generalizability from 3-T to 7-T fMRI and from resting-state to task-based fMRI. Trained on 3-T rs-fMRI data, the VAE was directly applied to 7-T rs-fMRI data without fine-tuning. The model encoded and compressed 7-T fMRI spatial patterns as vectors embedded in a 256-dimensional latent space and used these latent representations to reconstruct the input patterns. As shown in Figure 3, reconstruction yielded high squared spatial correlations (*r*^2^ = 0.71 *±* 0.02) with the input patterns spatially smoothed with FWHM=6 mm. This reconstruction accuracy for 7-T rs-fMRI data even surpassed that for 3-T rs-fMRI data (*r*^2^ = 0.68 *±* 0.02), likely due to the higher signal-to-noise ratio (SNR) given the higher field strength.

**Figure 3:**
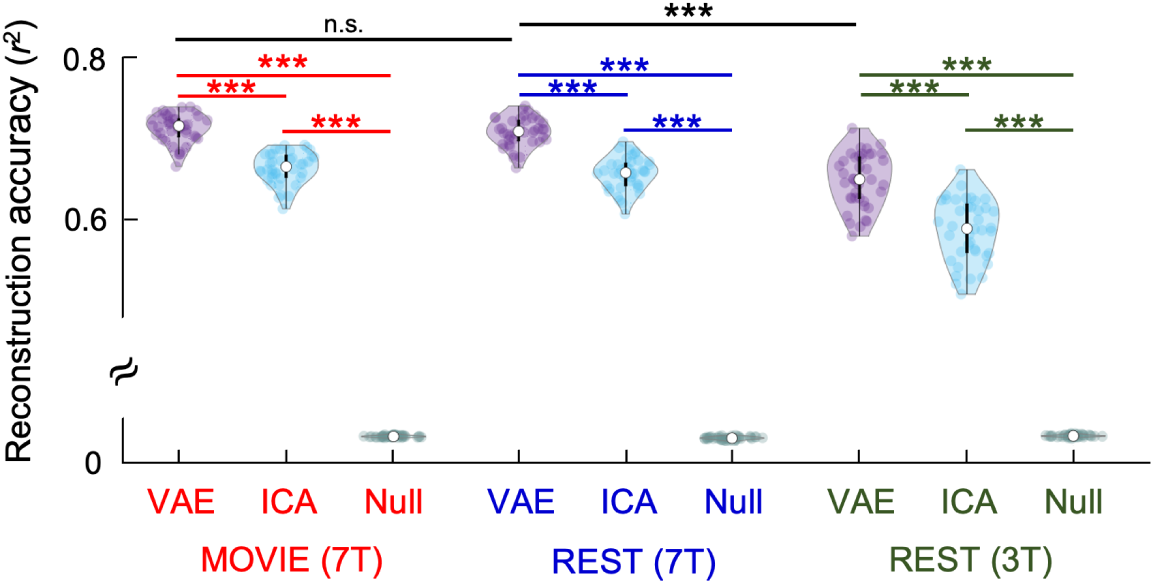
Model generalizability. Two models (VAE and ICA) pretrained with 3-T rs-fMRI data were tested for their performance in compressing unseen fMRI data into 256 latent variables or independent components and reconstructing the data from the compressed representations. As a baseline (Null), we also used the VAE model to generate fMRI activity patterns from random samples drawn from the prior latent distribution (i.e. spherical Gaussian distribution). The model performance, in terms of the squared spatial correlation (*r*^2^) between the reconstruction and the input spatially smoothed with FWHM = 6 mm, is evaluated for three datasets: 3-T rs-fMRI (green), 7-T rs-fMRI (blue), and 7-T fMRI during movie watching (red). The difference between models or datasets is tested for statistical significance using two-sample t-tests (***: *p <* 0.001 and n.s.: not significant).

To assess generalizability to task states, we applied the model to 7-T fMRI data during movie watching. Reconstruction accuracy under this task state was comparable to that at rest (no significant difference, *p* = 0.2). Across all datasets, the VAE consistently outperformed its linear counterpart, ICA, in compressing and reconstructing fMRI data (*p <* 0.001, paired t-test) (Figure 3). These results demonstrate that the VAE generalizes well across fMRI field strengths and brain states, extending beyond the conditions it was trained on.

### Latent Distributions of Resting-State vs. Movie-Evoked Activity

We next compared the latent distribution of 7-T fMRI during rest and movie-watching. At rest, the latent distribution exhibited a spherical geometry consistent with the prior distribution imposed during training (i.e., independent and standard Gaussian distributions). The covariance matrix of this distribution, averaged across subjects, approximated an identity matrix (Figure 4), indicating that rs-fMRI activity engaged all latent dimensions equally. Different dimensions corresponded to spatially overlapping resting-state networks that exhibit spontaneous cortical activity with non-linearly independent dynamics.

**Figure 4:**
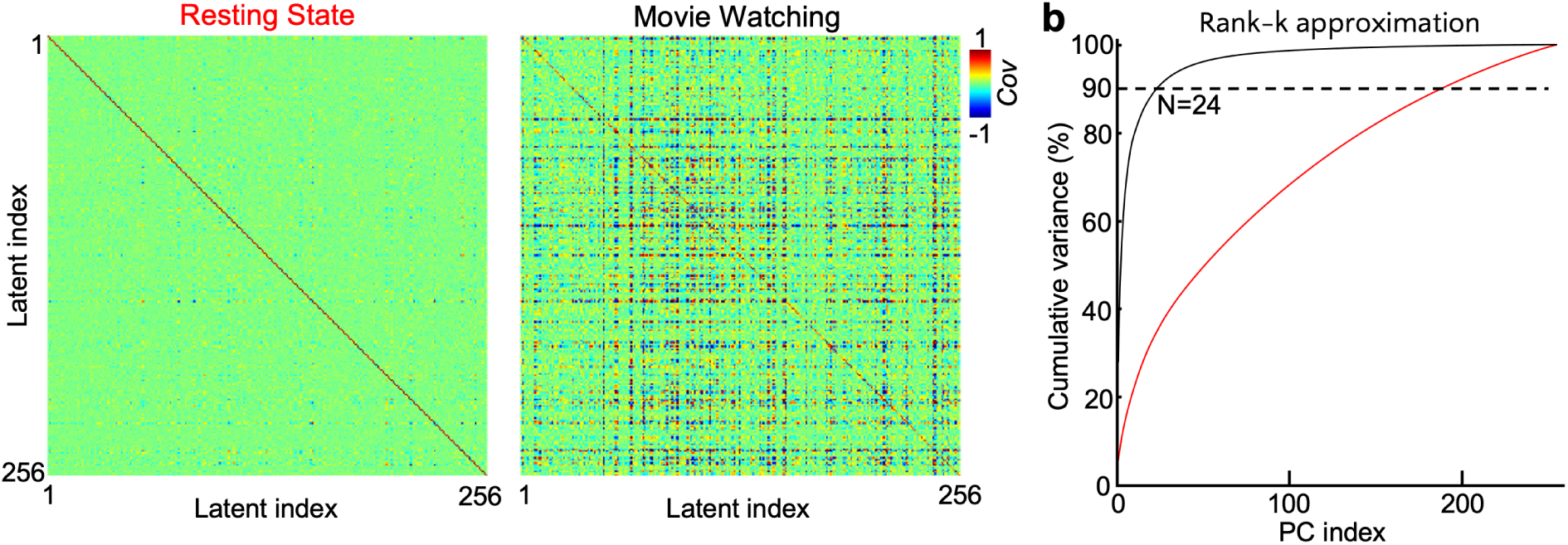
Latent representation geometry for spontaneous or movie-evoked activity. **a**. The representation geometry was characterized by the covariance matrix of the latent distribution for the resting state (left) or during movie watching (right). Cov: Covariance. The range of covariance is rescaled: mean variance (i.e., mean of diagonal elements of the covariance matrix) equals to 1. **b** shows the variance explained by the rank-k approximation of the covariance matrix for the resting state (red) or movie watching state (black), where *k* is the number of principal components that spanned a *k*-dimensional subspace. To explain 90% of the variance, the resting state required *k* = 189, and the movie watching state required *k* = 24.

In contrast, during movie-watching, the latent distribution deviated from a spherical geometry. The movie-evoked responses exhibited non-zero covariance between distinct latent dimensions (Figure 4b). A lower-rank approximation of the movie-evoked covariance matrix revealed that 24 dimensions explained 90% of the total variance, compared to 189 dimensions required for the same explanatory power at rest (Figure 4b). This suggests that brain activity driven by naturalistic stimuli engages about 13% of the intrinsic dimension of the resting state networks encoded by the model.

### Variation across Individuals and between Brain States

We examined how individual differences and brain states differentially influenced the latent representations and dynamics of cortical activity. Specifically, we analyzed the transitions in cortical activity over a 30-second interval, focusing on shifts from resting (0–15 seconds; right before Northwest) to movie watching (16–30 seconds; after Northwest). The t-SNE visualization of latent representations in a 2-D space revealed a clear and robust transition between resting and movie-watching states, consistent across 173 subjects (Figure 5, left). The large discontinuity between each subject’s resting-state and movie-watching representations suggests that transitions between brain states dominate over individual differences. However, subject latent representations remain separated within each brain state (Figure 5, right).

**Figure 5:**
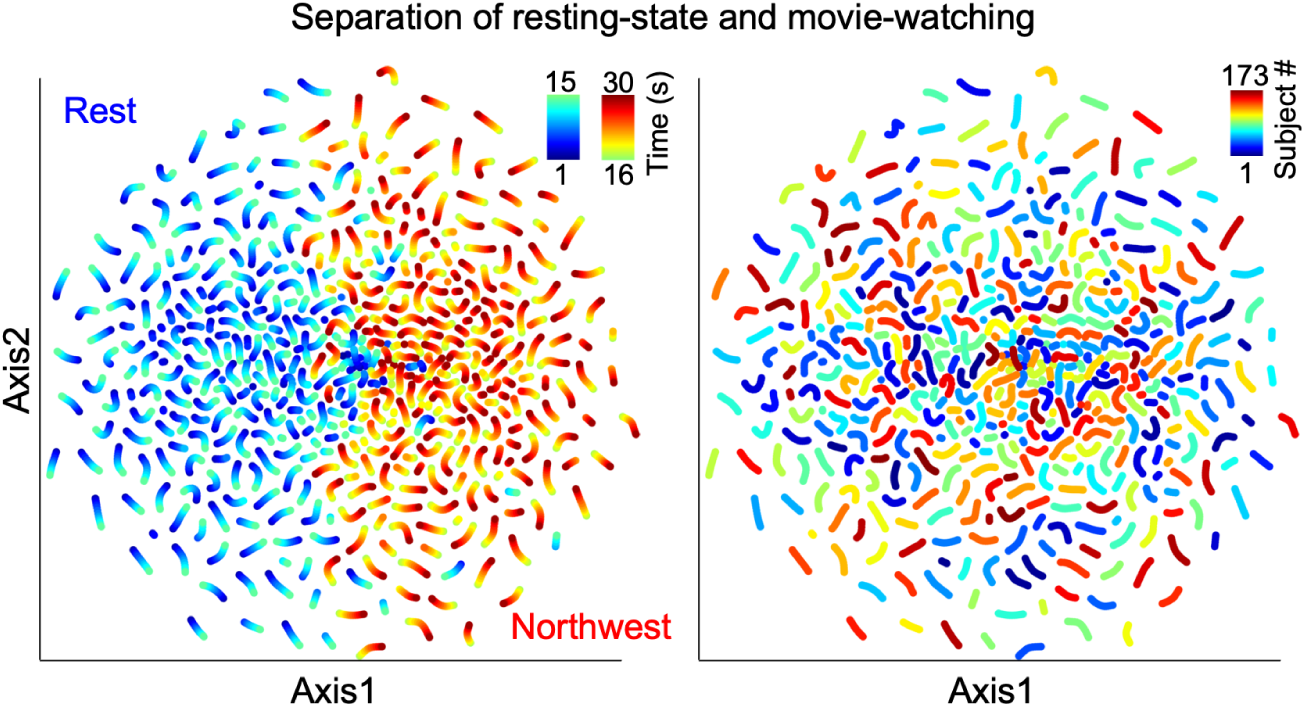
Variation in latent-space trajectory across brain states and individuals. The t-SNE visualization depicts latent representations of cortical activity over a 30-second interval as 173 subjects transitioned from resting (1–15 seconds) to watching a movie (“Northwest”) (16–30 seconds). Representations are color-coded by time (left) and by subject (right). The visualization demonstrates that latent representations are primarily influenced by brain state transitions (resting-state vs. movie-watching) rather than individual differences.

### Cortical Dynamics in the Latent Space

Extending this analysis to longer periods (*>* 60 minutes across 4 sessions) of alternating different movies and resting periods, we observed an evolving trajectory in the latent space. The velocity of this trajectory quantified the direction and magnitude of changes in brain activity over time (Figure 6). The time series of the latent velocity reveals abrupt changes, likely indicating discrete brain events occurring during movie watching.

**Figure 6:**
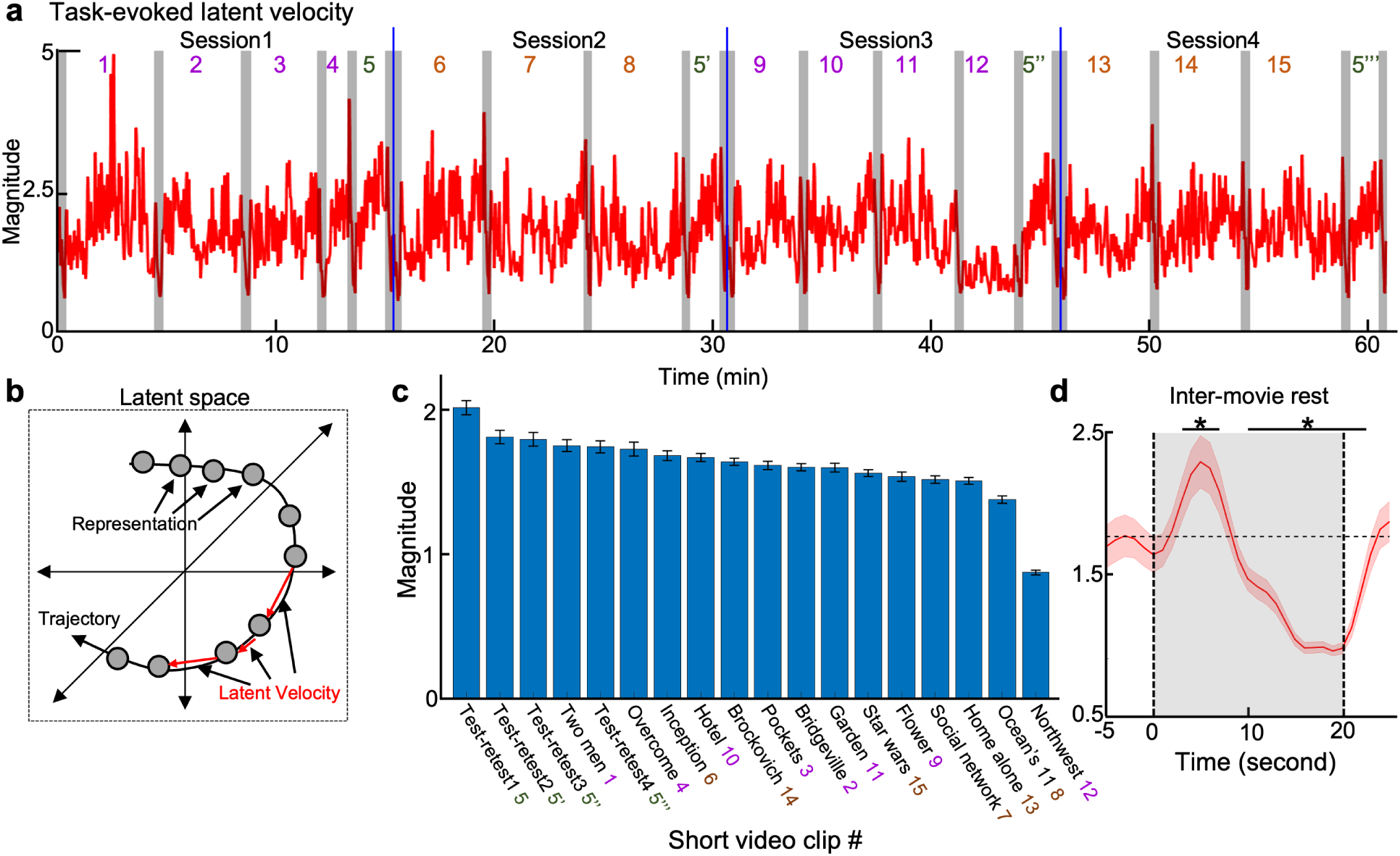
Latent velocity of cortical activity averaged across subjects reveals neural events during movie-watching and quantifies varying levels of brain engagement across movie clips. **a**. The magnitude of the latent velocity is shown as a time series (red) concatenated across four sessions (sessions 1–4), during which subjects alternated between resting periods and watching various movies (movies 1–15). Note that the movie 5 (“Test-retest”) was repeated four times (5, 5*^′^*, 5*^′′^*, 5*^′′′^*). Resting periods between sessions are marked in blue, while resting periods between movies are shown in gray. **b**. As latent representations (circles) evolve over time, their instantaneous changes (red arrows) define the latent velocity, representing the velocity of the latent trajectory. **c**. The average magnitudes of the latent velocity during movie watching are compared across different movies in descending order. **d**. Changes in the latent velocity magnitude from movie-watching to resting periods are averaged across all transitions, capturing the cortex-wide response to brain-state shifts. On top of this response, the periods of statistical significance (one-sample t-test, *p <* 0.05, marked as *) are shown as horizontal lines.

The magnitudes of the latent velocity differed significantly across different movies (ANOVA, *F* = 39.96, *p <* 10*^−^*^4^). This variability appeared to reflect the level of cortical engagement elicited by each movie (Figure 6c). For instance, the video “Northwest”, which mainly showed natural scenes (without any human-related features such as faces), had much lower cortical engagement than other movies that were more visually or emotionally stimulating. Moreover, greater latent velocity was observed during the initial watching compared to subsequent re-watching of the same clips, with the latent velocity for the test-retest clip greater for 5 than 5*^′^*, which is greater than 5*^′′^*, which in turn is greater than 5*^′′′^* (Figure 6c). This indicates possible neural adaptation processes due to repeated experiences. Short breaks (resting period) between movies triggered event-like responses in the latent space. During these breaks, the latent velocity increased rapidly, peaking at 5 seconds, decreasing to baseline by 10 seconds, and undershooting by 20 seconds (Figure 6d). These temporal characteristics align with the canonical hemodynamic response function, suggesting the neurogenic origin of the response observed in the latent space.

### Consistency across Sessions and Individuals

We analyzed a single movie clip (“Test-retest”, 84 seconds, repeated four times) to investigate how video content and features influenced the trajectory of latent representations. Given the first viewing of this movie, we examined the temporal alignment between video frames (Figure 7a) and latent representations visualized in 2-D using t-SNE (Figure 7b). The latent trajectory exhibited dynamic variability, showing intervals of gradual progression interspersed with abrupt transitions at specific time points (e.g., TR=4, 21, 48, 50, 61, 66, and 67). These transitions corresponded to notable changes in the movie’s audiovisual content, such as the onset of audio output at TR=4 and scene transitions or cuts at TR=21, 48, 50, 61, 66, and 67.

**Figure 7:**
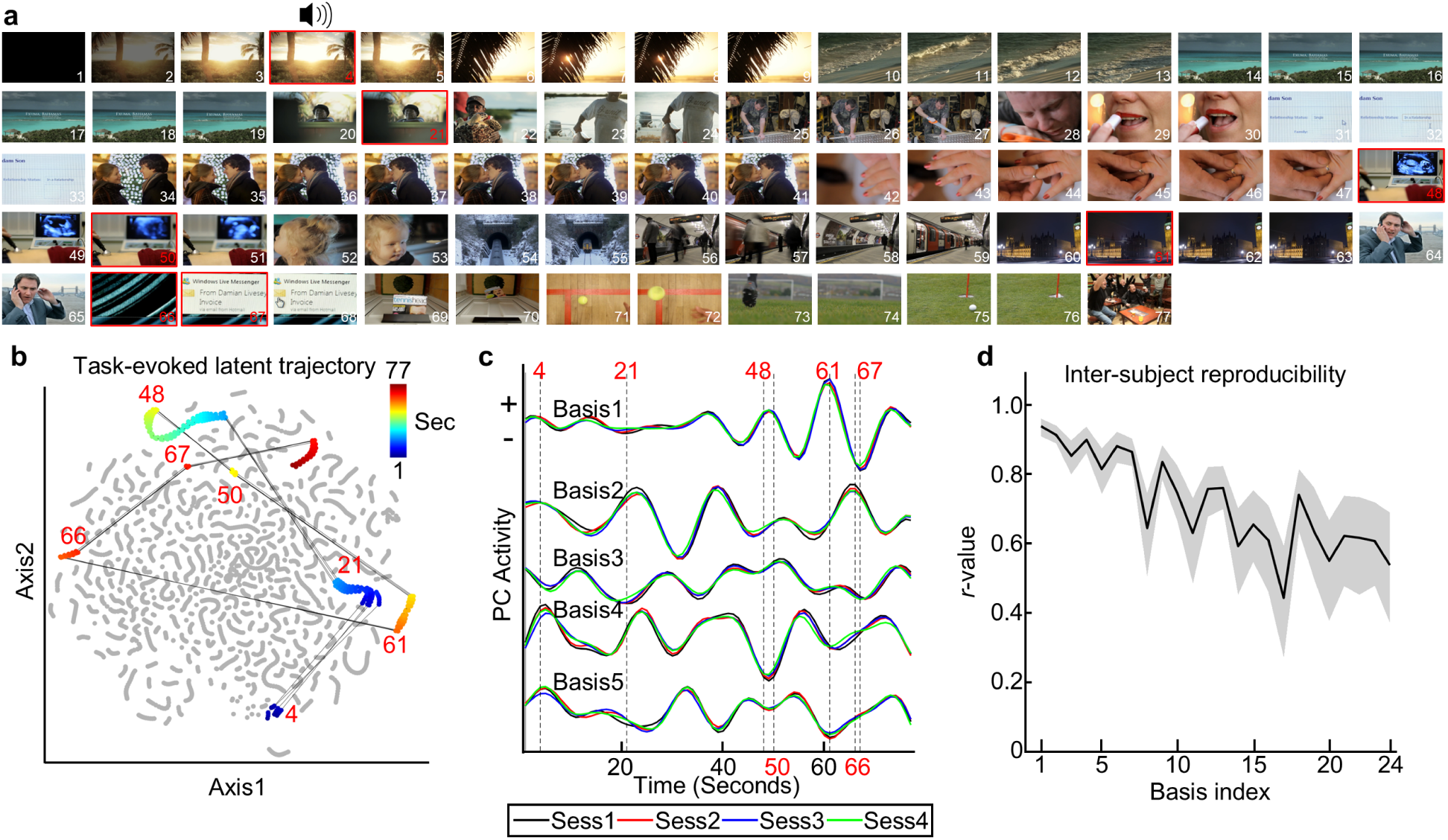
Latent trajectory of cortical responses to naturalistic movie stimuli. **a**. Selected video frames from the “Test-retest” movie, with their time of occurrence (in seconds) shown in the lower right corner in each frame. Frames corresponding to abrupt transitions in the latent trajectory are highlighted with red borders. The speaker icon indicates the onset of auditory stimuli. **b**. Latent representations visualized in 2D using t-SNE, applied to cortical activity from all individuals across all movies (the same 2D space as Figure 5). The latent trajectory during the “Test-retest” movie are highlighted in color, with colors indicating time within the movie. Straight lines represent abrupt shifts in latent representations, with red numbers indicating the time point of every shift. **c**. The top 5 principal components of the latent trajectory averaged across subjects during the “Test-retest” movie, along with their temporal dynamics for four repetitions (sessions 1-4, shown in different colors), demonstrating reproducibility across sessions. **d**. Correlations between group-averaged and individual-specific principal components for the top 24 components, demonstrating consistency across individuals. Upper and lower boundaries of shade stand for lower and upper quartiles, respectively. Line stands for the average *r*-score.

The latent trajectory was highly reproducible across the four repetitions of the “Test-retest” movie and consistent across individuals. To quantify this reproducibility, we averaged the latent trajectory across subjects for each session and extracted its principal components, representing the key dimensions of the latent space activated by this movie. The temporal dynamics of these principal components were highly consistent across all four repetitions (Figure 7c), indicating that the latent trajectory encoded distinct stimulus-locked temporal dynamics specific to the movie. Furthermore, these principal components exhibited high cross correlations across individuals (Figure 7d; mean of *r* = 0.93, 0.91, 0.85, 0.90, and 0.82 for basis 1–5, respectively; averaged after converting *r*-score to Fisher’s *z*-score), reflecting robust group-level cortical responses reproducible across individual subjects.

These findings demonstrate that the VAE-derived latent trajectories reliably capture stimulus-driven cortical responses and align closely with audiovisual events in naturalistic movie stimuli. The strong reproducibility across repeated sessions and individual subjects highlights the generalizability of this approach for characterizing group-level stimulus-evoked cortical responses.

## Discussion

In summary, this study supports the hypothesis that the cortical manifold learned from spontaneous activity constitutes a foundational state space for brain dynamics. We demonstrated the robust generalizability of our VAE model, trained on 3-T rs-fMRI data, across field strengths (3-T and 7-T) and brain states (resting and movie-watching). The latent representations confirmed that naturalistic movie stimuli are confined to a subspace in the latent space, occupying 13% of the functional reservoir defined by the resting state. Additionally, trajectories along this manifold were time-locked to audiovisual events, exhibited consistent dynamics across individuals, and showed high test-retest reliability. Our findings highlight the potential for using VAE-derived manifold geometry to characterize stimulus-driven cortical responses in dynamic and naturalistic contexts.

### VAE trained on resting-state fMRI generalizes to task conditions

Variability in fMRI data may reflect differences in acquisitions, paradigms, and subjects, etc. While training separate models for each condition is impractical, it is desirable to pretrain a model generalizable across many conditions. This requires large training datasets that capture diverse brain activity patterns. Rs-fMRI data are well suited for this purpose, as they encompass co-activation patterns consistent with those evoked by various tasks (Smith et al., 2009), making the training inclusive and unbiased. It is also desirable to train the model using unsupervised or self-supervised learning without being constrained or biased by any specific goals. Using unsupervised learning, the VAE is trained to compress inputs into latent representations and then use the compressed representations to reconstruct the input (Kingma & Welling, 2013; Higgins et al., 2017). This data-driven approach uses an information bottleneck for the model to learn how to extract essential information while minimizing redundancy by enforcing independence among latent dimensions.

Our results confirm the model’s generalizability without any fine-tuning. The VAE trained on 3-T rs-fMRI data could effectively encode and reconstruct 7-T resting-state data, demonstrating generalizability across field strengths. Our observation that the model showed better reconstruction accuracy on 7-T resting-state data than on 3-T training data is likely due to the higher SNR at higher field strengths and different parameters (e.g., a longer TR and a lower bandwidth) (Uğurbil et al., 2013). This improvement does not imply superior performance on out-of-distribution data. The model also generalized well to fMRI data during movie-watching. This extends the model’s applicability to diverse tasks, since watching a movie is a naturalistic condition involving complex sensory, emotional, and cognitive processing (Zhang et al., 2021). Even though movie-watching activity occupied only 13% of the resting-state–trained latent dimensions (24 vs. 189 PCs for 90% variance), it still achieved reconstruction accuracy comparable to resting-state data, indicating that robust stimulus-evoked BOLD patterns can be captured within a compact, low-dimensional subset of the intrinsic latent reservoir.

### Tasks engage distinct latent subspaces

The latent space learned from rs-fMRI data encapsulates a comprehensive reservoir of cortical networks. This reservoir reflects the intrinsic cortical functional architecture (Smith et al., 2009; Cole et al., 2014) grounded on underlying connectivity or geometry (Honey et al., 2009; Pang et al., 2023). Specific tasks engage only subsets of this full reservoir, activating cortical responses that are embedded within distinct latent subspaces. For instance, naturalistic movie stimuli activated approximately 13% of the entire latent space, as demonstrated in this study.

Mapping task-based functions in the latent space can provide a novel analysis framework for fMRI. Unlike traditional methods that map functions onto discrete brain voxels or regions (Friston et al., 1994), using independent latent variables allows for greater statistical rigor and facilitates algebraic operations to interrogate brain functions. For example, we may parcellate the latent space into task-specific subspaces. Different tasks may correspond to distinct latent subspaces, which may be overlapping, complementary, or nested within each other. Characterizing these relationships can inform how brain functions cluster into domains and subdomains (Insel et al., 2010; Poldrack et al., 2011; Laird et al., 2011). We can also compare latent representations during unknown tasks with those associated with known tasks. By assessing the affinity or distance to these subspaces, we may decode brain activity into specific tasks or combinations of tasks.

### Latent trajectory quantifies cortical dynamics and engagement

The trajectory of the latent representation characterizes cortical dynamics. Variations in the velocity of the latent trajectory may reflect changes in brain states (e.g., transitions from rest to movie watching) or external inputs (e.g., audio onset, scene shifts). The latent dynamics are consistent with the temporal characteristics of the established neurovascular model (Boynton et al., 1996; Lindquist et al., 2009), indicating their neurogenic origin. Furthermore, the average velocity of the latent trajectory during movie watching is consistent for the same movie but varies reliably across different movies. This measure provides a quantifiable index of cortical engagement elicited by a movie, potentially serving as a ”brain score” to rate movies based on their ability to elicit cortical dynamics. Such a score may potentially complement subjective popularity ratings (e.g., IMDb), offering neural evidence-based insights for movie production and evaluation.

### Individual variations and commonality

Every brain is unique, and individual variations shape the functional connectivity in the resting state (Finn et al., 2015; Gratton et al., 2018). The VAE-based latent representations of resting state activity preserve and even enhance individual differences, enabling reliable individual identification (Kim et al., 2021). However, during movie watching or transitions from rest to tasks, nearly all individuals exhibit consistent changes in the latent trajectory. This suggests that common group factors dominate over individual variations in shaping the latent dynamics of task-based fMRI activity, unlike those of resting state activity. This is consistent with prior findings that naturalistic stimuli elicit neural responses similar across individuals (Hasson et al., 2004). However, it is possible that naturalistic stimuli also enhance individual differences in terms of emotional and cognitive responses (Finn & Bandettini, 2021).

Future studies should further characterize and differentiate individual variations and commonality in the latent space. Characterizing individual variations is critical to personalized medicine for mental illness or neurological disorders, where individual-specific deviations from common patterns may provide diagnostic insights (Marquand et al., 2016). Conversely, characterizing individual commonality is essential to neuroscience for understanding brain functions at a population level (Glasser et al., 2016).

## Conclusion

This study establishes that the intrinsic manifold learned from spontaneous activity serves as a universal scaffold for cortical dynamics, and constrains cortical responses to the environment. Our VAE model pretrained on rs-fMRI data generalizes effectively across field strengths, brain states, and naturalistic stimuli. The model can capture the latent trajectory of cortical dynamics and reveal variation across individuals and brain states.

## Acknowledgment

The research is supported by the National Institutes of Health MH104402, AG082204, National Science Foundation IIS 2112773, and the University of Michigan.

## Code Availability

The code and weights for the VAE model are available at https://github.com/libilab/rsfMRI-VAE.

---

**Table S1:** Information sheet of video clips used in the study.

| Sess. | Short title | Original title (year) | Src | TR | Short description | Note |
| --- | --- | --- | --- | --- | --- | --- |
| 1 (1) | Two men | Two Men (2009) | Vimeo | 245 | A man sees two men running and talk about possible motives | With subtitle and narration |
| 2 (1) | Bridgeville | Welcome to Bridgeville (2011) | Vimeo | 221 | People explain why they love their small town |  |
| 3 (1) | Pockets | Pockets (2008) | Vimeo | 189 | People explain their items in pocket and significance of items |  |
| 4 (1) | Overcome | Inside the Human Body (2011) | Vimeo | 65 | Short documentary about people overcome physical disabilities | With narration |
| 5 | Test-retest | 23 Degrees South, LXIV (2011) | Vimeo | 84 | Concatenation of short videos (2-6 s) covering various objects |  |
| 6 (2) | Inception | Inception (2010) | Hwd | 228 | A man introduce how one can manipulate another person's dream | Metaphysical e.g., Moving building |
| 7 (2) | Social Network | The social network (2010) | Hwd | 260 | Mark Zuckerberg's disciplinary hearing and its aftermath |  |
| 8 (2) | Ocean's 11 | Ocean's Eleven (2001) | Hwd | 251 | A man explains the plot heisting a casino with visual aids |  |
| 5' | Test-retest |  |  |  |  |  |
| 9 (3) | Flower | Off The Shelf (2008) | Vimeo | 181 | A journey of a flower from the house to the field (with music) | Only music |
| 10 (3) | Hotel | 1212 | Vimeo | 185 | Metaphysical encounter between a man and woman in a hotel room |  |
| 11 (3) | Garden | Mrs. Meyer's Clean (2013) | Vimeo | 205 | Documentary about a woman who runs an urban garden community |  |
| 12 (3) | Northwest | Northwest Passage | Vimeo | 143 | Portray of wet landscapes and abandoned buildings | No human presented |
| 5" | Test-retest |  |  |  |  |  |
| 13 (4) | Home alone | Home alone (1990) | Hwd | 234 | A kid is searching his home, realizing he is alone at home | Poor video quality |
| 14 (4) | Brockovich | Erin Brockovich (2000) | Hwd | 231 | A woman talks with a woman and visits a legal office with her children |  |
| 15 (4) | Star wars | The Empire Strikes Back (1980) | Hwd | 257 | A scene on an icy planet and Han Solo visit headquarter of rebellion |  |
| 5''' | Test-retest |  |  |  |  |  |

